# Natural hybridisation reduces vulnerability to climate change

**DOI:** 10.1101/2022.10.25.513775

**Authors:** Chris J. Brauer, Jonathan Sandoval-Castillo, Katie Gates, Michael Hammer, Peter J. Unmack, Louis Bernatchez, Luciano B. Beheregaray

## Abstract

Understanding how species can respond to climate change is a major global challenge. Species unable to track their niche via range shifts are largely reliant on genetic variation to adapt and persist. Genomic vulnerability predictions are used to identify populations that lack the necessary variation, particularly at climate relevant genes. However, hybridization as a source of novel adaptive variation is typically ignored in genomic vulnerability studies. We estimated environmental niche models and genomic vulnerability for closely related species of rainbowfish (*Melanotaenia* spp.) across an elevational gradient in the Australian wet tropics. Hybrid populations between a widespread generalist and narrow range endemics exhibited reduced vulnerability to projected climates compared to pure narrow endemics. Overlaps between introgressed and adaptive genomic regions were consistent with a signal of adaptive introgression. Our findings highlight the often-underappreciated conservation value of hybrid populations and indicate that adaptive introgression may contribute to evolutionary rescue of species with narrow environmental ranges.

## Main

The environmental conditions experienced by species throughout their evolutionary history contribute to determine their present-day niche and constrain their distributional limits ^1,2^. Ancestral environments also shape contemporary patterns of standing genetic variation that are a key component of evolutionary potential. On the one hand, species that evolved in narrow environmental ranges may lack variation at genomic regions important for adaptation to a changing environment. On the other hand, generalist species that tolerate a much wider range of conditions may be better placed to respond to rapid climate change.

In predicting species’ vulnerability to rapid climate change, three evolutionary responses are typically considered; genetic adaptation, dispersal to a more suitable environment, or acclimation to the altered environment through phenotypic plasticity ^3^. An alternative and perhaps complementary evolutionary mechanism that is less often assessed is inter-specific introgression following hybridisation. That is the transfer of genetic material from one species into another by repeated backcrossing. Through this process, vulnerable species may adopt and exploit aspects of the evolutionary history of species more suited to the changed environmental conditions ^4,5^.

The role of hybrid populations in conservation is controversial due to concerns about diluting the genetic integrity of parental species, as well as policy and legislative uncertainty ^6^. Hybridisation can potentially increase the risk of extinction via outbreeding depression, through demographic swamping due to infertile or maladapted hybrids, or by genetic swamping leading to complete replacement of the local gene pool ^7,8^. The threat posed by these issues however is likely case-specific and may be less of a concern if hybridisation is natural and has occurred over an extended period. Introgression as a source of novel genetic variation that can increase evolutionary potential is only recently gaining widespread appreciation, particularly in animals ^9^. This has led to a call for hybrid populations to be given greater conservation value in policy and management decisions ^10^. Hybrid zones could potentially facilitate evolutionary rescue of many species threatened by climate change ^5^. This concept is also the basis of proposals for human-mediated evolutionary rescue of threatened species via translocations ^4,11,12^.

Genomic vulnerability assessments are increasingly used to identify populations that lack genetic variation likely to be important for adaptation to climate change ^13,14,15^. A range of statistical methods have been employed ^16,17,18,19^, however the basic framework is mostly similar regardless of the approach (but see ^20^). The first of two steps is to build a statistical model of the relationship between putatively adaptive genetic variation and the current environment. Secondly, this model is applied to projections of future environmental conditions to predict the change in allele frequencies required to maintain present patterns of local adaptation (also termed genomic offset). In addition to estimating the amount of evolutionary change required, it is equally important to understand the capacity for that change to occur naturally. This second component, rarely assessed in studies of genomic vulnerability, infers whether the adaptive alleles are present in a population. Most studies have focussed on abiotic factors influencing vulnerability, however species interactions and other evolutionary processes can also impact species’ responses to climate change ^21,22,23^.

Here we explored an ideal biogeographic scenario involving rainbowfishes endemic to the Wet Tropics bioregion of north-eastern Australia to assess if natural hybridization can influence vulnerability to climate change. Rainbowfishes (genus *Melanotaenia*) are a suitable ectotherm system to investigate climatic-driven adaptive evolution ^24,25,26,27,28,29,30,31^. They are a species-rich group of small fish found across the full spectrum of freshwater habitats in the Australian continent ^32^. Their adaptive capacity to respond to projected climates appear to be biogeographically determined ^27^ and their patterns of local adaptation are linked to hydroclimatic gradients and divergent thermal environments ^26,28,29,30^. When experimentally exposed to future climates, rainbowfishes show associations between upper thermal tolerance and gene expression responses influenced by the biogeographic context where species evolved ^25,27,31^.

They also display range-wide differences in genotype and environment associations linked to seasonal variation in stream flow and temperature ^28,29^, as well as to phenotypic traits that affect fitness ^26,30^. The combined evidence from experimental and wild populations of *Melanotaenia* species ^24,25,26,27,28,29,30,31^ support the hypothesis that historical climatic variation, regional climatic differences, and population connectivity influence variation in rainbowfish traits that determine regional patterns of adaptive resilience to climate change.

In this study we targeted five closely related species of tropical rainbowfishes that differ in predicted sensitivity to climate change. These include a widespread lowland generalist, *Melanotaenia splendida splendida*, and four narrow range specialists; the upland species *M. eachamensis*, Malanda rainbowfish and Tully rainbowfish (the latter two are undescribed), and the lowland species, *M. utcheensis*. We henceforth refer to the four narrow range species as narrow endemic rainbowfishes (NERs). *Melanotaenia splendida* are widely distributed and abundant across north-eastern Australia, whereas the NERs exhibit restricted distributions confined to short river valleys within and below the Atherton tablelands (Fig. 1A) ^33^. The study area is centred at a well described contact zone between lineages of many species expanding from two major Quaternary refugia ^34,35,36,37,38,39^. The Australian Wet Tropics bioregion is listed as a World Heritage Area and as a biodiversity hotspot. It is severely threatened by climate change, with the extinction of many endemic species predicted as temperatures increase and the cooler upland rainforest habitat disappears ^40^. Since cessation of volcanic activity in the early Holocene, major drainage patterns have largely resembled the present-day arrangement ^41^ and it is expected that *M. splendida* and the NERs have intermittently been in contact during most of that time. In the absence of large waterfalls separating populations, species boundaries (and potentially hybrid zones) have most likely been maintained by local hydroclimatic conditions associated with each species climatic niche. In the last few decades, climate change has resulted in *M. splendida* encroaching further into higher elevation habitat occupied by the NERs, and hybrids have been found where the species meet ^42,43^. This has raised concerns over the potential for NER populations to become threatened with extinction due to hybridisation with *M. splendida* ^43^.

**Figure 1.**
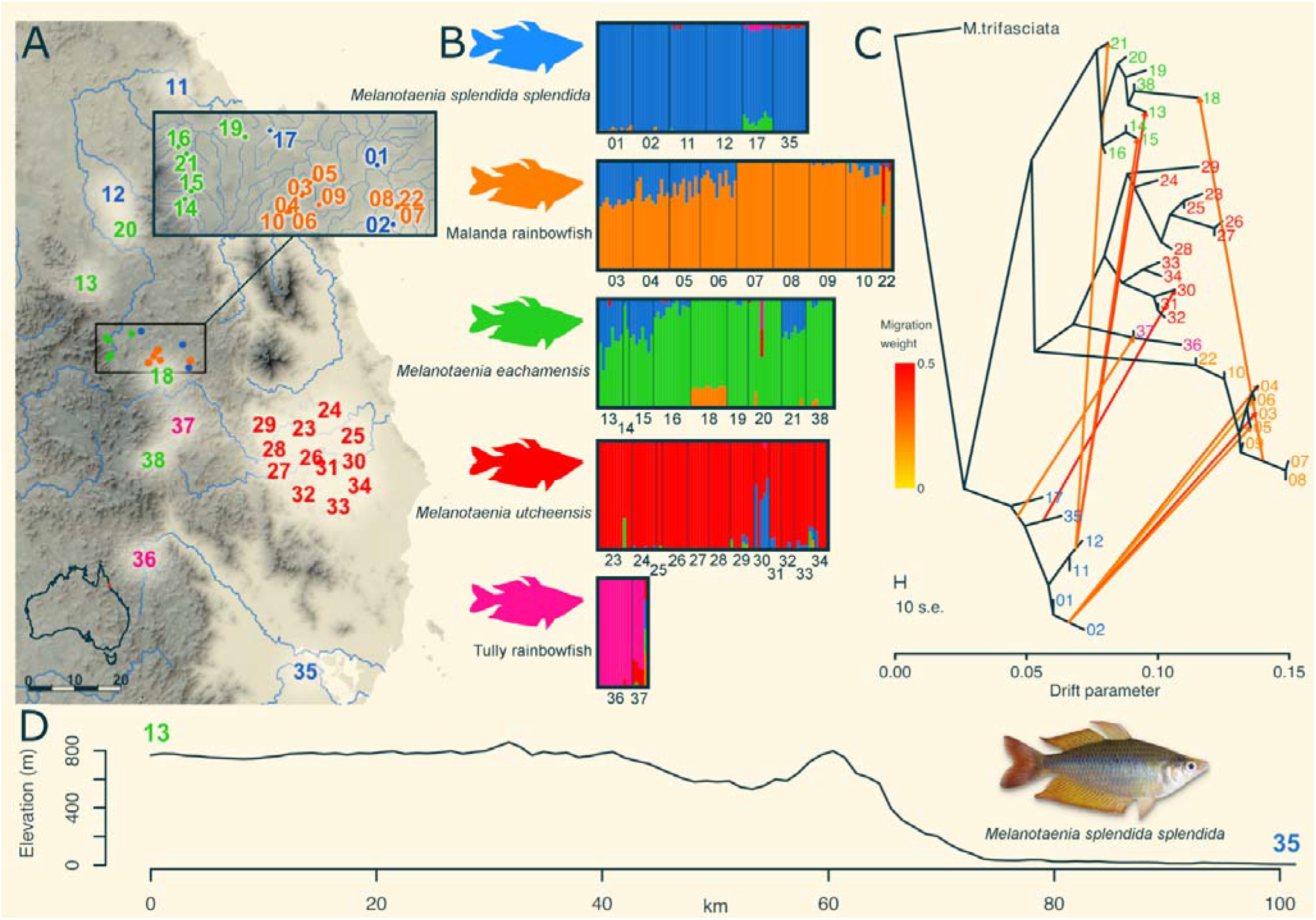
Sampling locations and spatial patterns of hybridisation for *Melanotaenia splendida*, Malanda rainbowfish, *M. eachamensis*, *M. utcheensis* and Tully rainbowfish. A) Sampling sites in the Wet Tropics of Queensland, Australia. B) Admixture plots for K=5 ancestral species. C) Treemix maximum likelihood tree showing introgression among species. D) Topographic relief profile indicating the difference in elevation between upland and lowland habitat along a transect between sites 13 and 35.

We constructed environmental niche models (ENMs) for all species to track range size variation throughout the Holocene and into the future. We predict that NERs, but not *M. splendida*, will lose large areas of suitable habitat under projected climates. We then used genotype-environment association analyses to identify candidate adaptive variation to estimate genomic vulnerability. We predict that hybrid NER populations will retain more adaptive alleles in the future and will require less evolutionary change to maintain patterns of local adaptation than pure NER populations. We also extended the genomic vulnerability framework to infer historical evolutionary responses to variation in climate throughout the Holocene. That extension enables that estimates of future vulnerability are interpreted in the context of the rate and magnitude of past environmental changes. By examining patterns of hybridisation and introgression between *M. splendida* and the NERs, we hypothesise that introgressive hybridization could be a source of novel adaptive variation likely to facilitate evolutionary rescue of species threatened by climate change.

### Genomic variation, hybrid detection and introgression

We examined 13,734 single nucleotide polymorphism (SNP) loci to reveal extensive patterns of hybridisation and introgression among the 344 individuals representing the five species (Fig. 1; Supplementary Table 1). More than 98% of SNPs mapped to one of 24 pseudo-chromosomes and were evenly distributed (mean = 565, SD = 68.7 SNPs per chromosome; Supplementary Table 2). Estimates of genetic diversity varied across species and among sampling sites but were significantly elevated (pairwise Wilcoxon rank sum <0.01) for hybrid populations compared to pure NER populations (Supplementary Tables 3-4).

Individual ancestry proportions ^44^ for K=5 ancestral species (Fig. 1B) were used to classify 167 individuals as pure (Q>0.95), and 85 individuals as *M. splendida*-NER hybrids (*M. splendida* and one NER Q>0.1, all other species Q<0.05). Pure individuals included 41 *M. splendida*, 22 *M. eachamensis*, 36 Malanda rainbowfish, 57 *M. utcheensis* and 11 Tully rainbowfish. We found hybrids between *M. splendida* and *M. eachamensis* (31), Malanda rainbowfish (49) and *M.utcheensis* (5), while no hybrid Tully rainbowfish were identified. Hybrid index ^45^ estimations among *M. splendida* and each NER supported the pure and hybrid classifications (Extended Data Fig. 1). Parental interspecific heterozygosity was marginally greater than expected based on hybrid indexes, suggesting the presence of ancestral polymorphisms in both parental species. Hybrid individuals showed reduced levels of interspecific heterozygosity (Extended Data Fig. 2-4), providing evidence for advanced-generation hybrids. Our ML tree demonstrated that each lineage is monophyletic, however the best model supported nine migration events from *M. splendida* into NERs and one between NER species (Fig. 1C). NewHybrids ^46^ simulations demonstrated the high power of our data to resolve hybrid classes providing an overall posterior probability >0.995 across all simulated genealogical classes for each species. Empirical results supported the other analyses in identifying advanced generation hybrids for all species (Supplementary Table 5.; Extended data Fig. 1-2). Overall, our findings support the hypothesis that contact between the narrow endemics and *M. splendida* has likely been recurrent over long periods of time.

To assess historical admixture and genome-wide introgression, we calculated *D*, *f*_4_-ratios and *f*_dM_ statistics for all *M. splendida*-NER trios with Dsuite. We found strong evidence for introgression between *M. splendida* and *M. eachamensis*, *M. utcheensis* and Malanda rainbowfish with a significant excess of ABBA (positive *D*) for all trios (Table 1). Positive *f*_4_-ratios were also observed for all trios, indicating substantial introgression from *M. splendida* into the three NERs (Table 1). Results for the sliding window statistic *f*_dM_ identified 28 introgressed regions for the hybrid individuals from *M. eachamensis*, 26 for the hybrid Malanda rainbowfish, and 18 introgressed regions for *M. utcheensis*. The introgressed regions were well distributed across 23 of 24 pseudo-chromosomes (Supplementary Table 2) and contained 153, 89 and 61 SNPs mapping to the main dataset for *M. eachamensis,* Malanda rainbowfish and *M. utcheensis*, respectively.

**Table 1.** Evidence for introgression from *M. splendida* to *M. eachamensis* and Malanda rainbowfish using D and *f*_4_-ratio statistics calculated by Dsuite. BBAA are derived alleles shared by (P1,P2), ABBA are derived alleles shared by (P2,P3). BABA are derived alleles shared by (P1,P3).

| P1 | P2 | P3 | D | Z-score | $f_4$-ratio | BBAA | ABBA | BABA |
| --- | --- | --- | --- | --- | --- | --- | --- | --- |
| Malanda | hybrids | *M. splendida* | 0.435 | 25.63 | 0.28 | 1017.31 | 305.34 | 120.31 |
| *M. eachamensis* | hybrids | *M. splendida* | 0.277 | 20.41 | 0.32 | 901.11 | 393.93 | 223.24 |
| *M. utcheensis* | hybrids | *M. splendida* | 0.236 | 12.84 | 0.25 | 766.96 | 394.28 | 243.78 |

### Environmental niche models (ENMs)

The ENMs based on maximum temperature of the warmest month (Bio05) and precipitation of the coldest quarter (Bio19) for *M. splendida*, *M. eachamensis*, *M. utcheensis* and Malanda rainbowfish (Extended Data Fig. 3) provided a good fit for the current species distributions (Supplementary Table 6). Throughout the Holocene, suitable habitat area for the NERs remained stable and similar to their recent historical known range. However, Malanda rainbowfish, for example, are predicted to lose as much as 92% of their current range by 2070 under the intermediate (RCP4.5) emissions scenario and more than 95% under the high (RCP8.5) emissions scenario (Fig. 2). A large area of high habitat suitability was confirmed for *M. splendida* throughout the Holocene, and ENMs based on projected 2070 climate suggested that most of that area is likely to remain suitable for the species (Fig. 2). Due to the low number of occurrence records for Tully rainbowfish (they are only known from four locations), ENMs could not be estimated for this species.

**Figure 2.**
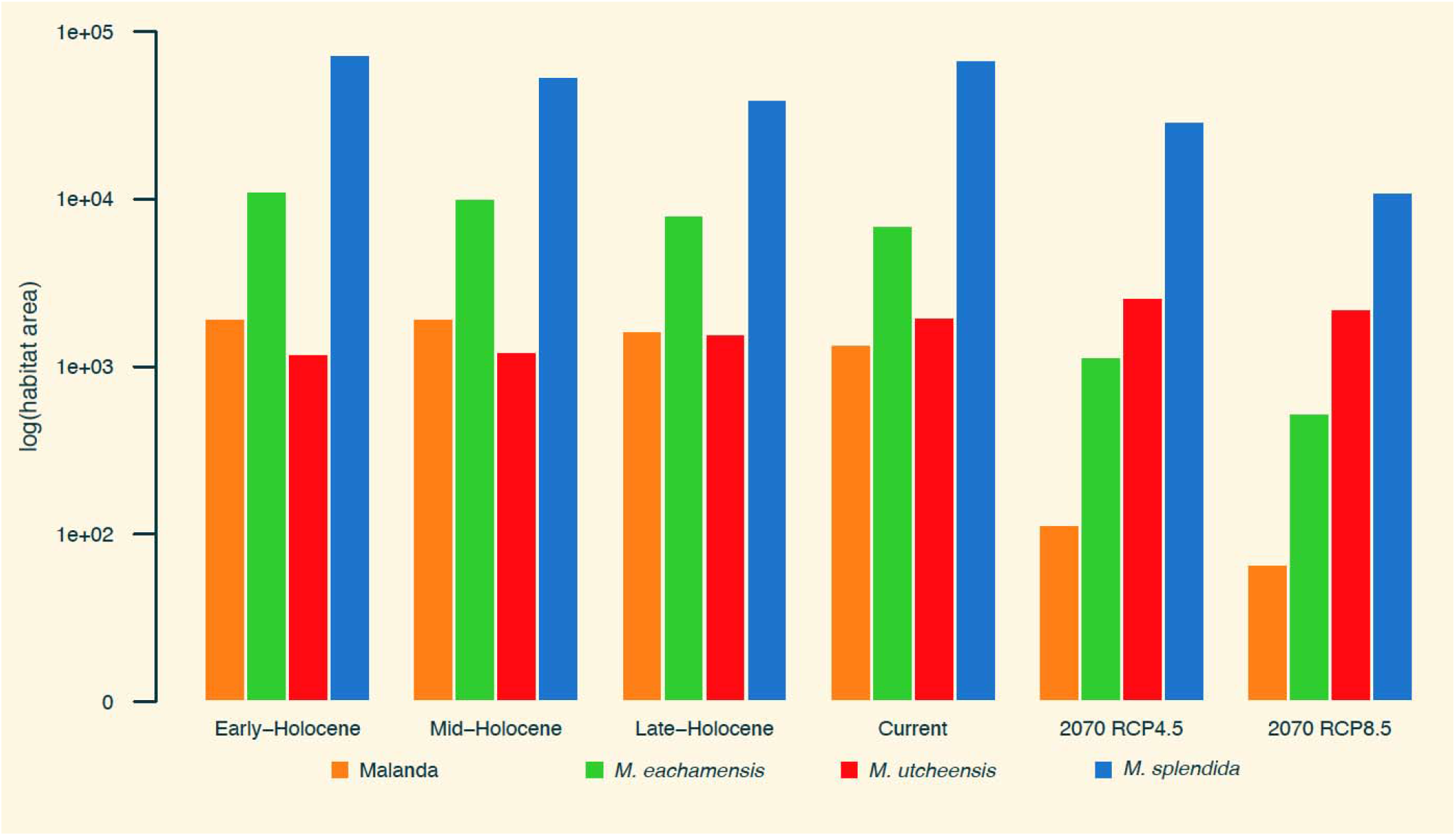
Reduction in suitable habitat area inferred for rainbowfish from the Wet Tropics throughout the Holocene and predicted for 2070. Narrow endemic species, Malanda rainbowfish, are predicted to lose up to 95% of their current distribution by 2070 (RCP8.5 emissions scenario). Habitat area is expressed as the log number of suitable pixels on a binary representation of ensemble environmental niche models using a probability of occurrence threshold of 70% to estimate range sizes at each time period.

### Climate adaptation and genomic vulnerability

The genotype-environment association analysis, temporal climatic change models and genomic vulnerability analyses together demonstrate the threat climate change poses to NERs (Fig. 3). Maximum temperature of the warmest month (Bio05) and precipitation of the coldest quarter (Bio19) were retained for the GEA analysis after accounting for correlated variables. The RDA model was significant (*R*^2^=0.15, *P*<0.001) and identified 211 candidate adaptive SNPs associated with the two climatic variables, with the first two axes explaining 15.99% and 12.84% of the variation constrained by the environment (Fig. 3A; Supplementary Tables 7-8). Four candidate adaptive SNPs occurred in introgressed regions identified using the *f*_dM_ sliding window statistic. These included one for Malanda–*M. splendida* hybrids and three for *M. eachamensis*–*M. splendida* hybrids. In total 163 GEA candidate SNPs were annotated to 158 genes, and 294 introgressed candidates were annotated to 201 genes (Supplementary Table 9). Gene ontology enrichment analyses revealed several enriched categories for the GEA candidates (q<0.05), including mitogen-activated protein kinase (MAPK) signaling pathway genes that have regulatory functions on heat shock proteins and cellular responses to thermal stress ^47^ (Supplementary Table 10). Annotation of genomic positions and predicted functional effects indicated that a substantial number of candidate SNPs were in exons, with expected moderate to high impact on gene function (Extended data Fig. 4).

**Figure 3.**
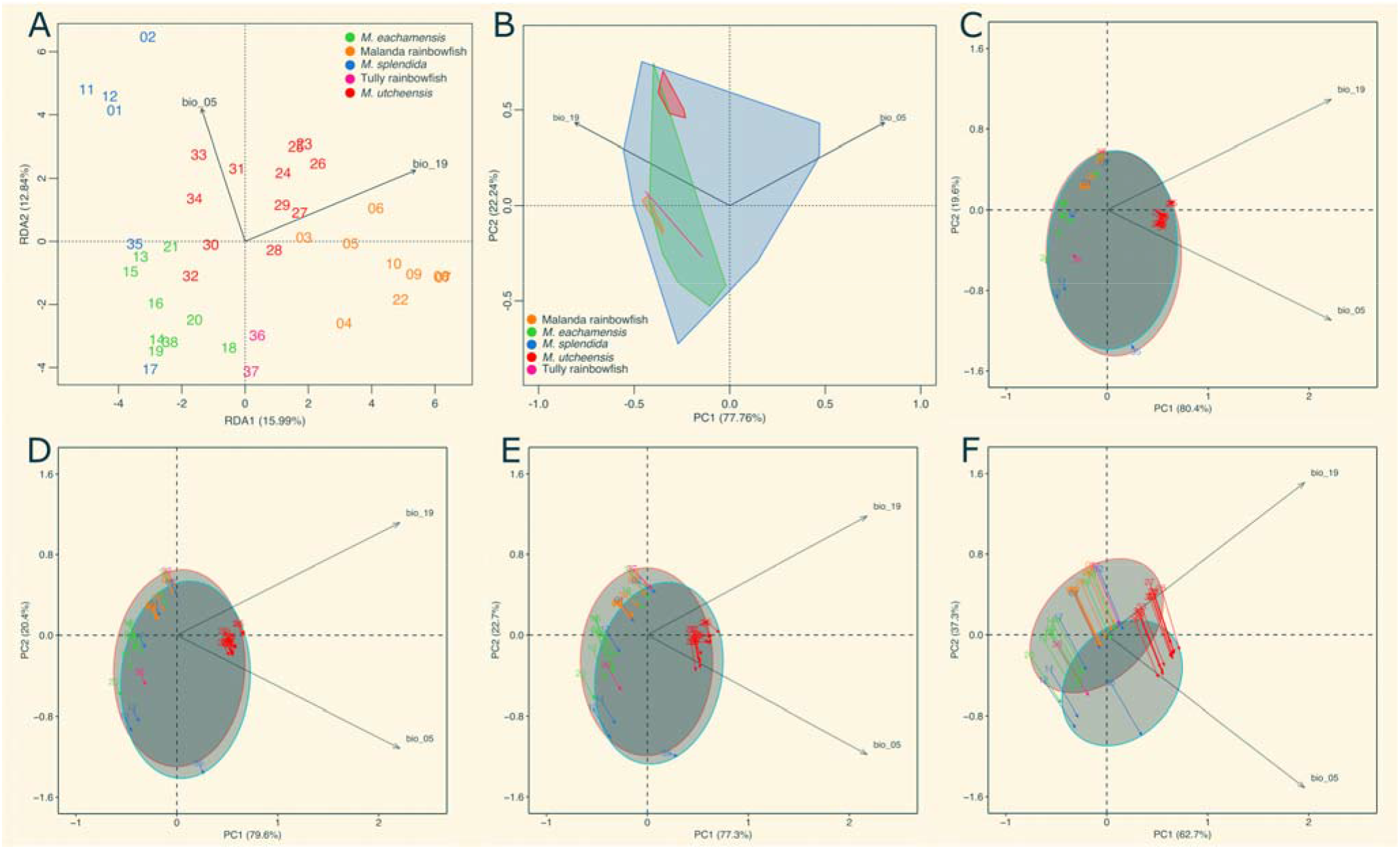
Genotype–environment association and principal component analyses (PCA) reveal potentially adaptive genomic variation, the relative size and position of the current climatic niche for each species, and modelled changes in climate between the early Holocene and 2070 at each sampling site. **A)** Biplot summarising the redundancy analysis. **B)** Current climate PCA indicating the relative size and position of each species niche. Temporal PCA plots demonstrating direction and magnitude of climate change between time periods; **C)** early Holocene to mid Holocene, **D)** mid Holocene to late Holocene, **E)** late Holocene to current, and **F)** current to 2070 (RCP8.5 projections).

The environmental niche PCA highlights the much larger environmental envelope occupied by *M. splendida* compared to NERs. The lowland specialist *M. utcheensis* inhabits a warmer and wetter environment than the upland species that are restricted to much cooler conditions (Fig. 3B). The climate change PCAs reveal relatively little variation in climate over approximately 10ky from the early Holocene until the present. However, for 2070 the climate is expected to become much hotter and slightly drier (RCP8.5 projections; Fig. 3C-F).

Following model calibration based on the current environment and observed candidate adaptive allele frequencies, eight populations for which the model performed poorly (*R*^2^<0.5) were not considered in the final vulnerability estimates ^48^. Allele frequencies were then predicted for the remaining 28 sampling sites for the historic (Fig. 4A) and projected future environments (Fig. 4B). *Melanotaenia splendida* ancestry was a reasonable predictor of genomic vulnerability (*R*^2^=0.11, *P*<0.04) supporting the hypothesis that introgression with *M. splendida* may provide an evolutionary rescue effect for NER species most vulnerable to climate change (Fig. 4C). Allele frequencies of the candidate loci were estimated for each species and the percentage of loci missing the adaptive allele (based on the future climate models) were aggregated for pure and hybrid populations (Fig. 4D). Populations of pure Malanda rainbowfish missed 41% of adaptive alleles, whereas hybrid Malanda populations had variation at >99% of candidate loci. For *M. utcheensis*, 27% of adaptive alleles were missing from pure populations with just 2% of adaptive alleles absent from hybrid populations. Pure populations of *M. eachamensis* appeared less depauperate than the other NERs with 12% of adaptive alleles absent, while only 7% of adaptive variation was missing from hybrid populations. In contrast, pure *M. splendida* populations were missing just 0.5% of adaptive alleles.

**Figure 4.**
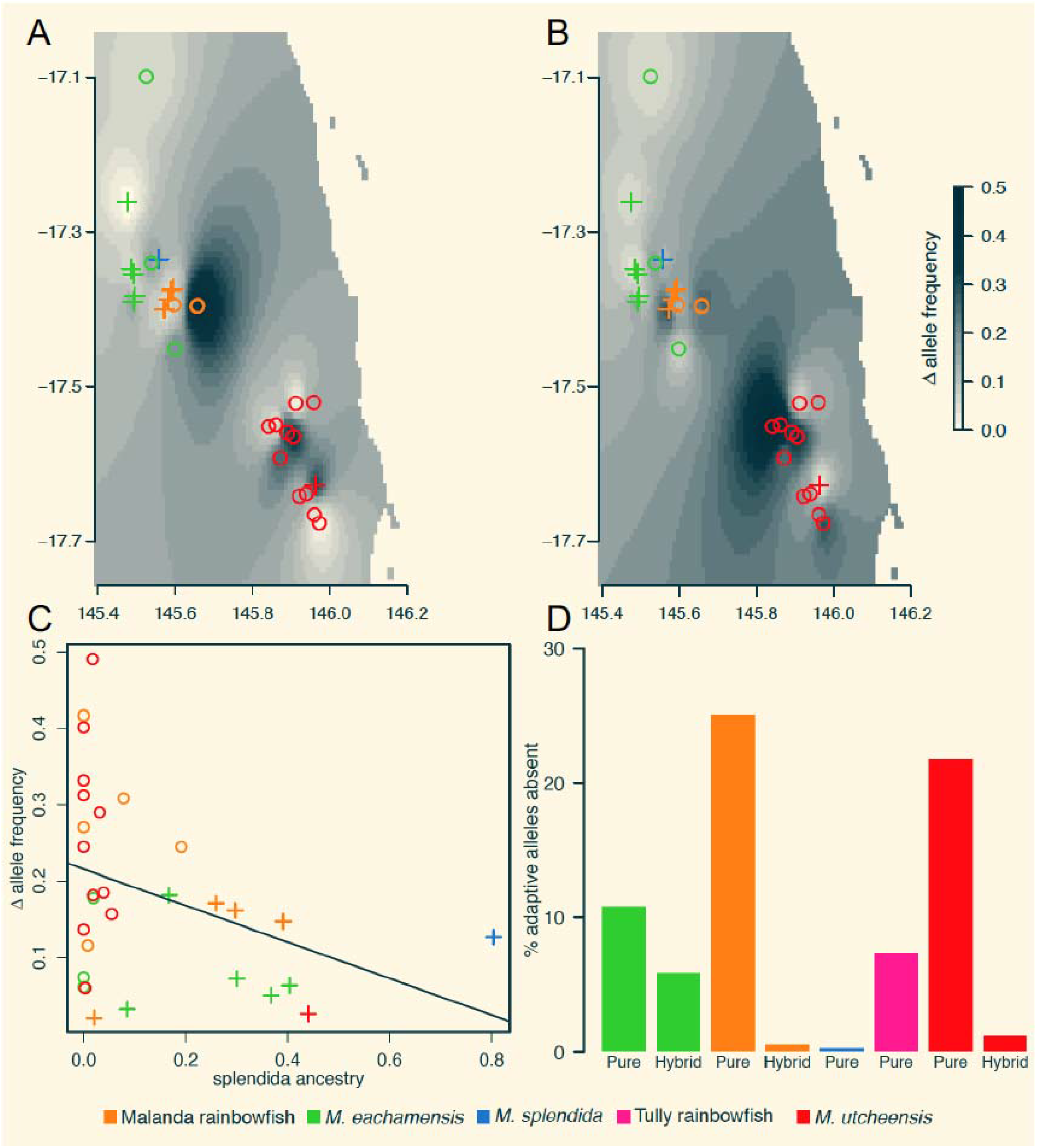
Genomic vulnerability is reduced for hybrid populations of rainbowfish. Estimates of the allele frequency change required to keep pace with climate change for *Melanotaenia eachamensis*, Malanda rainbowfish, Tully rainbowfish, and *M. utcheensis* based on the candidate adaptive SNPs for **A)** late-Holocene to current and **B)** current to 2070 (RCP8.5). **C)** required change in allele frequency as a function of *M. splendida* ancestry (*R*^2^=0.11, *P*<0.04). **D)** Percentage of adaptive alleles (relative to 2070 RCP8.5 climate predictions) absent from pure, and hybrid populations. Hollow circles – pure populations, crosses – hybrid populations.

## Discussion

Our genomic vulnerability assessments reveal that populations of narrow endemic rainbowfishes that demonstrate introgressive hybridisation with a warm-adapted widespread generalist also exhibit reduced genomic vulnerability to climate change. This supports the hypothesis that natural evolutionary rescue may moderate the effects of climate change for these populations. Examining the genomic vulnerability estimates in the context of historical environmental niche models (ENMs) indicated that the evolutionary change required in the next 50 years far exceeds that which has likely occurred since the early Holocene. These findings are consistent with evidence from experiments of adaptive resilience to projected climates and from range-wide surveys of adaptation that indicated that physiological performance limits and adaptive capacity in rainbowfishes are closely linked to local climatic conditions and range sizes ^25,26,27,28,29,30,31^. Our approach expands on assessments of genomic vulnerability ^15,17,49,50,51^ by considering the potential for adaptive introgression to enhance longer-term genomic responses to rapid environmental changes. Hybrid populations were shown to be less vulnerable, based on both the amount of evolutionary change required (i.e. adaptive allele frequency change), and the capacity for that change to occur naturally (i.e. whether the adaptive alleles are present). The latter is an often-overlooked component of genomic vulnerability, and one that highlights the importance of standing genetic diversity for evolutionary potential.

Increasing empirical work demonstrates that introgression can accelerate adaptive shifts in response to environmental change ^52,53,54,55^ that might help lineages threatened by climate warming ^56,57,58^. For example, there is strong evidence for adaptive introgression between archaic humans ^59^, and also among more modern human populations ^60^ that likely facilitated rapid adaptation to new environments. Nolte et al. ^61^ identified a hybrid population of sculpins (*Cottus gobio*) that were able to invade habitat unsuitable for either of the two parental lineages. One striking aspect of this example is the speed at which the hybrid population was able to adapt, with evidence that divergent phenotypic and life history traits arose along with the new habitat preferences in just a few decades. Our findings are consistent with a signal of adaptive introgression that could promote evolutionary rescue of cool-adapted species. This interpretation however requires further validation via common garden experiments (e.g. ^62^) to provide fitness estimates for the parental and hybrid lineages in current and predicted temperatures.

Contact between the cool adapted rainforest species and warm adapted lowland species has likely occurred recurrently throughout the Quaternary, consistent with findings for many lineages in this region ^63,64^. Rainbowfish ecotypes are known to be constrained by climate ^26,27,28,29,30^, and previous experiments of upper thermal tolerance and adaptive resilience to projected temperatures indicate that biogeographic factors might strongly influence climate change vulnerability ^25,27,31^. The hybrid zones examined here have likely also been maintained by the respective climatic niches of the parental species, however the ENMs predict that by 2070 Malanda rainbowfish may lose >95% of suitable habitat, and *M. eachamensis* face >92% reduction (RCP8.5 projections). As the cooler upland climatic niche retreats, it is unclear to what extent pure and hybrid NER populations might either persist or be replaced completely by *M. splendida*. Translocating pure populations of upland species outside of their current range is unlikely to provide a long-term solution as they currently occupy the only remaining cool tropical montane rainforest region on mainland Australia. Their impending niche loss coupled with high genomic vulnerability potentially provides few options for conservation managers in the future. We suggest that populations with low levels of introgression (e.g. sites 10, 16) should not be cause for concern and should be afforded the equivalent conservation status as pure populations. We also argue that more advanced hybrid populations should be conferred greater value for their potentially crucial role in retaining unique diversity from the NER lineages in the future. These populations may also buffer species-level vulnerability in the short-term, either directly via introgressed adaptive alleles, or through indirect effects such as increased local effective population sizes ^65^. Interestingly, although *M. utcheensis* is a lowland species and has evolved in a warmer environment, populations of this species exhibit some of the highest genomic vulnerability and lowest adaptive capacity to future conditions. While the coastal lowland environment of *M. utcheensis* appears to have remained relatively stable throughout the Holocene, populations may have been isolated by their unique climatic niche and we found little evidence for gene flow via hybridisation. This highlights that the evolutionary history of specialist warm-adapted species could render them more vulnerable to climate change than might be expected.

In 1985, Soule ^66^ described the (then) emerging field of conservation biology as a crisis discipline, and this has never been more true than now. The rate of anthropogenic climate change is challenging many species to mount evolutionary responses to environmental changes occurring on an ecological time scale. Conservation biologists and managers are also increasingly obliged to make difficult decisions without the time or resources necessary to fully understand the potential implications of these decisions. The genetic and demographic consequences of hybridisation are difficult to predict but should be carefully considered when assessing whether possible negative effects are offset by potential gains in adaptive resilience. Here we identified long-established hybrid rainbowfish populations harbouring potentially important and novel genetic variation for responding to climate change. These populations would typically be ignored in management plans that focus on maintaining pure lineages ^67,68^. The high-resolution of modern genomic techniques can reveal subtle signals of admixture and we must be cautious in how we interpret invasiveness and the threat of hybridisation ^69^. Nonetheless, the ability to more precisely characterise ancestry means that patterns of hybridisation can be well defined. Our work highlights the conservation value of hybrid populations and exemplifies how adaptive introgression may contribute to natural evolutionary rescue of species threatened by climate change.

## Online Methods

### Study system

Thought to have an old origin, rainbowfishes (Melanotaeniidae) are the most speciose freshwater fish family endemic to Australia and New Guinea ^70^. Multiple lineages exist sympatrically which occasionally hybridise ^32,70^, and many species readily do so in captivity ^71^. This tendency may have helped facilitate their rapid adaptive radiation (e.g. ^72^) across the diverse range of climatic ecotypes they now inhabit in Australia ^32^. Previous work, including for the generalist *M. splendida splendida*, has identified genomic signatures of local adaptation and adaptive plasticity associated with biogeographic history, hydroclimatic variation, and projected climates ^25,26,27,28,29,30,31^. Additionally, rainbowfishes are renowned for their high morphological diversity ^73,74^, and strong links between morphological variation and local adaptation have also been established for several Australian species, including for the NER, *M. eachamensis* ^24,30,75^.

### Sampling and Genomic Data

Samples for 344 individuals from five Australian rainbowfish species ^70^ were collected from 38 sites in the Australian wet tropics (Table 1). In addition to the five focal species, two samples from a sixth rainbowfish species, *M. trifasciata* were also included as an outgroup ^70^. Fish were either sampled live and returned to the water with caudal fin-clips stored in 100% ethanol, or euthanized in an overdose of AQUI-S^®^ solution (50% isoeugenol), frozen in liquid nitrogen, and stored at −70°C in the Australian Biological Tissues Collection at the South Australian Museum, Adelaide.

DNA was extracted following a modified salting-out protocol (Sunnucks & Hales 1996) with DNA assessed for integrity using gel electrophoresis and for purity with a NanoDrop 1000 spectrophotometer (Thermo Scientific). Double digest restriction-site-associated DNA sequencing libraries ^76^ were prepared using the restriction enzymes *SbfI* and *MseI* (New England Biolabs). Using custom individual barcodes to multiplex samples (96 per lane for all ingroup samples and 48 per lane for the outgroup samples), libraries were randomly assigned to each of seven Illumina HiSeq2500 lanes and sequenced as single-end, 100-bp reads. Raw sequencing data were demultiplexed using the *process_radtags* module from STACKS 2.4 ^77^. Individual fastq files were trimmed using Trimmomatic v0.39 ^78^ and aligned to a *M. duboulayi* reference genome using Bowtie2 v2.3.5.1 ^79^. The SAM files were converted to BAM files and duplicate reads were marked and removed using Picard v2.21.7 (https://github.com/broadinstitute/picard), before using GATK v3.8-1-0 ^80^ for indel realignment. BCFtools v1.9 ^81^ (bcftools call -m) was used to call SNPs. Raw genotypes were filtered for missing data, mapping quality (>30), HWE, MAF (>0.01) before pruning to reduce the effect of linkage disequilibrium (LD). We first estimated LD decay across the genome and plotted pairwise R^2^ among 66,762 raw SNPs before fitting a spline of exponential decay to estimate the distance in base pairs at which decay is no longer significant (p>0.05) based on Tukey’s criteria for anomalies. We found that average R^2^ does not change significantly after 605bp and pruning SNPs <300 bp apart resulted in 99.3% of SNP pairs separated by >100Kbp, with <0.005% separated by less than 600bp (Supplementary Table 11).

### Genomic variation, hybrid detection and introgression

Genetic diversity summary statistics expected heterozygosity (He), observed heterozygosity (Ho) and percentage of polymorphic loci were estimated for each sampling site and for aggregated pure and hybrid populations per species using the hierfstat R package ^82^. We used a Wilcoxon rank sum test implemented in the stats R package ^83^ (pairwise.wilcox.test) to assess differences in heterozygosity among pure and hybrid populations of the NERs.

To identify individuals with hybrid ancestry we first used ADMIXTURE v1.3.0 to estimate individual ancestry proportions (Q) assuming a K value of five ancestral species ^44^. We determined pure individuals as those with a Q-value of >0.95. Hybrid status was assigned to individuals with both *M. splendida* and one other species ancestry >0.1, with the remaining species Q-values <0.05. This allowed the evaluation of introgression between *M. splendida* and each of the narrow endemic species while reducing noise associated with individuals with multiple species ancestry. To additionally assess patterns of hybrid ancestry we estimated hybrid indices using the method implemented in the gghybrid R package ^84^ and generated triangle plots to visualise the relationship between interspecific heterozygosity and hybrid index. We also performed simulations using NewHybrids v.1.1 ^46^ and the Hybriddetective R package ^85^ to test the power of our data for detecting hybrids and to assign individuals to hybrid classes. We selected panels of ~200 informative SNPs and generated three replicates of three simulations with pure parents, F1, F2, and backcrosses between F1 and pure parents. Samples were then assigned to hybrid classes, based on the posterior probability thresholds estimated with the simulations. We used a Jeffreys-like prior and default genotype proportions with a burn-in of 20,000 iterations followed by 200,000 MCMC sweeps.

We used Treemix ^86^ to examine introgression between branches of the rainbowfish phylogeny. This method models both topology and gene flow by first using allele frequencies and a Gaussian approximation for genetic drift to estimate a maximum likelihood (ML) tree. The residual fit of the ML tree is used to identify populations that are a poor fit to the tree, before migration edges are fitted between branches in stepwise iterations to maximise the likelihood. We ran Treemix testing 1-20 migration events, using blocks of 500 SNPs (-k 500), no sample size correction (-noss) and two *M.trifasciata* samples as an outgroup. The final model was selected as the number of migration events at the asymptote of the log-likelihood estimations for all models. Based on the ADMIXTURE results we used Dsuite ^87^ to assess gene flow between *M. splendida* and the other species and to identify introgressed loci. For these analyses we re-filtered the original raw genotypes, based on the reduced number of individuals (as described above). The data were again filtered for missing data (<20%), MAF (>0.01), however the HWE filter was not applied as divergent allele frequencies are expected among species. PLINK v1.9 ^88^ was used to prune the SNPs for linkage disequilibrium (--indep 50 5 2). The resulting 27,009 SNP dataset was used to calculated Patterson’s *D* ^89,90^, also known as the ABBA–BABA statistic based on the tree (((P1,P2),P3),O). The *D* and *f*_4_-ratio statistics were calculated using the Dtrios function in Dsuite with default parameters. Trios were assessed to test the hypothesis of introgression between *M. splendida* and each of the NERs for which hybrids fitting the above criteria existed. In this case we tested trios where P1 represented the pure narrow endemic samples, P2 the hybrid samples, P3 the pure *M. splendida* samples and O the outgroup, *M. trifasciata*. In addition to assessing evidence for gene flow between *M. splendida* and the narrow endemics, we also estimated the sliding window statistic *f*_dM_ ^91^ to identify specific introgressed genomic regions. Implemented with the Dinvestigate function in Dsuite, a sliding window of 50 SNPs with a step of 10 SNPs was used (-w 50,10). Windows in the top 5% of the *f*_dM_ distribution were considered as candidate introgressed loci. Overlapping candidate windows were merged using BEDtools v2.29.1 ^92^ to provide a minimum set of candidate regions for each trio. We then used BCFtools view to map the 13,734 SNP dataset to the candidate introgressed regions to identify any overlap with the candidate climate adapted SNPs.

### Ecological niche models

Bioclimatic (BIOCLIM) variables were extracted from CHELSA v1.1 ^93,94^. Projections for 2070 under intermediate (RCP4.5) and high (RCP8.5) emissions scenarios were also obtained from CHELSA based on the Australian Community Climate and Earth System Simulator (ACCESS1.0) global circulation model ^95^. Historic climate models (also derived from CHELSA) were downloaded from PaleoClim ^96^ for the early-Holocene (11.7-8.326 ka), mid-Holocene (8.326-4.2 ka) and late-Holocene (4.2-0.3 ka) ^97^. These data were resampled from 2.5 arc-minutes to match the current and future datasets 30 arc-second resolution. All rasters were cropped to an area encompassing the catchments from which samples were obtained. Coastlines for all time periods were also cropped to the current coastline to control for the effect of sea level changes throughout the Holocene. This was to enable direct comparison of habitat suitability across time periods for the specific extent of potential habitat available now and in the future (2070).

To predict species vulnerability to climate change, ecological niche models were generated for each species and each time period using biomod2 v3.4.6 ^98^. In addition to locations for the genomic samples, occurrence data for a further 420 locations within the study extent were obtained from the Atlas of Living Australia (ALA: http://www.ala.org.au). These data were filtered for duplicate entries, geographic accuracy and to remove outliers based on known distributional limits. To avoid collinearity among variables, and to reduce the likelihood of overfitting the RDA and environmental niche models, we initially conducted a PCA on raster data from all 19 BIOCLIM variables across the study area using the raster_pca function from the synoptReg R package ^99^. We then selected one temperature and one precipitation variable that most highly correlated with the first two axes of the initial climate PCA to ensure that spatial variation in climate was well captured. The ecological importance of the retained variables, maximum temperature of the warmest month (Bio05) and precipitation of the coldest quarter (Bio19), has also previously been demonstrated in studies of rainbowfish adaptation ^26,27,28^. Ensemble models were built using four commonly used algorithms Maximum Entropy (Maxent), Generalised Linear Model (GLM), Generalised Boosting Model (GBM), and Random Forest (RF) ^100^. Five hundred pseudo-absences were randomly selected from the model extent. Each model was replicated three times and those with a relative operating characteristic (ROC) curve statistic >0.8 were retained. The weighted mean of probabilities ensemble models for each species were converted to binary representation using a probability threshold of 70% and used to estimate relative range sizes at each time period. A more sophisticated method of determining the binary threshold (minimum suitability of the top 90% of training sites) was trialled initially, this was found to bias the narrow endemic species range estimates downwards due to limited occurrence records. After exploring a range of parameters, we found that using the 70% suitability threshold provided a good balance between model accuracy and precision across all species.

### Climate adaptation and genomic vulnerability

To identify a candidate set of climate adapted loci for Tropical Australian rainbowfish we employed a genotype–environment association (GEA) analysis using redundancy analysis (RDA) to detect associations between population allele frequencies and the same two climatic variables used for the ENMs (Bio05, and Bio19). To control for the non-linear spatial phylogenetic structure in the RDA, we estimated Moran’s eigenvector maps (MEM) ^101^ using the *mgQuick* function from the MEMGENE R package ^102^, before using a forward selection procedure to identify significant MEM eigenvectors to use as conditioning variables. We used the rda function in the vegan R package ^103^ and tested significance of the final model using the anova.cca function and 1000 permutations. The mean locus score across all SNPs was calculated for each of the first two RDA axes, and those scoring greater than three standard deviations from the mean were considered candidates for hydroclimatic selection ^104^. Overlap between the GEA and introgressed candidate loci identified were considered as potential signals of adaptive introgression. We used SnpEff ^105^ to perform gene, genomic position and functional effect annotations for the candidate loci based on the *M. duboulayi* genome. Gene ontology (GO) terms and Kyoto Encyclopedia of Genes and Genomes (KEGG) pathway enrichment analyses were explored using the STRING web server ^106^.

To assess changes in the environment since the early Holocene, and how climate is predicted to change over the next 50 years, principal components analyses (PCA) were performed on climate data for each time period based on the retained bioclim variables from the RDA. The population.shift function from the AlleleShift R package^19^ was then used to visualise and compare the magnitude and direction of environmental changes between periods. An additional PCA was also performed using the retained current environmental data and plotting convex hulls surrounding the sampling sites for each species to highlight the relative size and any overlap of the environmental niche space occupied by each species.

AlleleShift was also used to model the rate of past evolutionary change and to predict future genomic vulnerability based on the candidate adapted loci. An initial two step calibration first used RDA to build a model (AlleleShift::count.model) and predict the relationship between allele counts and the environmental data (AlleleShift::pred.model). Secondly, the predicted allele counts are used as independent variables in a generalised additive model with observed allele frequencies as the response (AlleleShift::freq.model). This step constrains the final allele frequency predictions to fall between 0 and 1. Model fit was then evaluated for each population and those for which the model performed poorly (*R*^2^<0.5) were omitted from the final analyses as suggested by Blumstein et al. ^48^. Based on the calibrated allele frequency–environment model, allele counts were predicted for the 2070 projected environmental data and then converted to allele frequencies to enable direct comparison. Genomic vulnerability was then expressed simply as the difference between median values of the observed and predicted allele frequencies among the current and projected environmental models (referred to as delta allele frequency. Allele frequency shifts were also estimated using the historic environmental models to help interpret the genomic vulnerability assessments in the context of inferred rates of evolutionary responses to climate change throughout the Holocene. To test the hypothesis that hybrid populations show reduced genomic vulnerability to climate change, we constructed a linear model examining the relationship between genomic vulnerability and the proportion of *M. splendida* ancestry for each population. Finally, to assess the capacity for the pure NER populations to adapt in situ assuming no gene flow from *M. splendida* or the hybrid populations, we identified for how many loci the adaptive allele (as predicted by the AlleleShift model) was missing.

## Author Contributions

C.J.B., J.S-C., K.G., P.J.U., L.B., and L.B.B conceived, designed, and refined the project. C.J.B. and J.S-C. performed the data analysis with assistance from K.G. and P.J.U. All authors contributed to interpreting the results. C.J.B. and L.B.B. led the writing of the manuscript, with input from J.S-C., K.G., M.H., P.J.U. and L.B.

## Acknowledgements

Collections were obtained under permits from various state fisheries agencies and research is under Flinders University Animal Welfare Committee approval E342. We thank Minami Sasaki for laboratory assistance. Financial support was provided by the Australian Research Council via grants to L. B. B. (FT130101068; DP150102903). Keith Martin provided samples, assistance in the field and valuable discussion about rainbowfish species distributions and habitat preferences.

## Data availability

The *Melanotaenia duboulayi* reference genome assembly is available at NCBI, accession number TBA. Genotypes for the 13,734 and 27,009 SNP datasets and environmental data files can be accessed on Dryad: doi:TBA.

## Code availability

Code can be accessed on GitHub at https://github.com/pygmyperch/NER

