## Extended data for "Natural hybridisation reduces vulnerability to climate change"

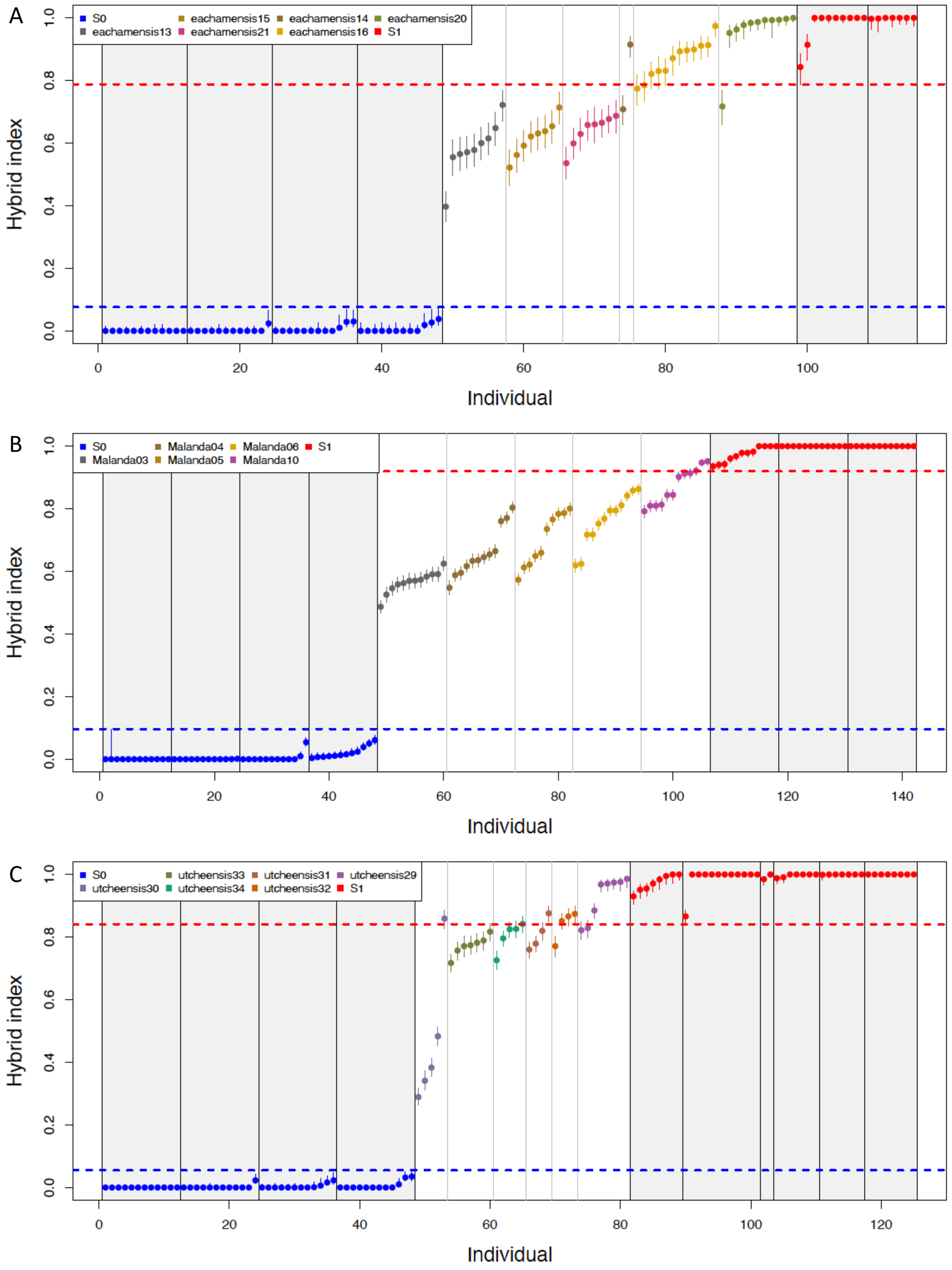


**Extended data Figure 1. Hybrid index estimations among *Melanotaenia splendida* and narrow endemic species (NERs) A) *M.eachamensis*, B) Malanda rainbowfish and C) *M. utcheensis* populations.** Pure reference populations are coded as S0 (*M. splendida*), S1 (NERs).

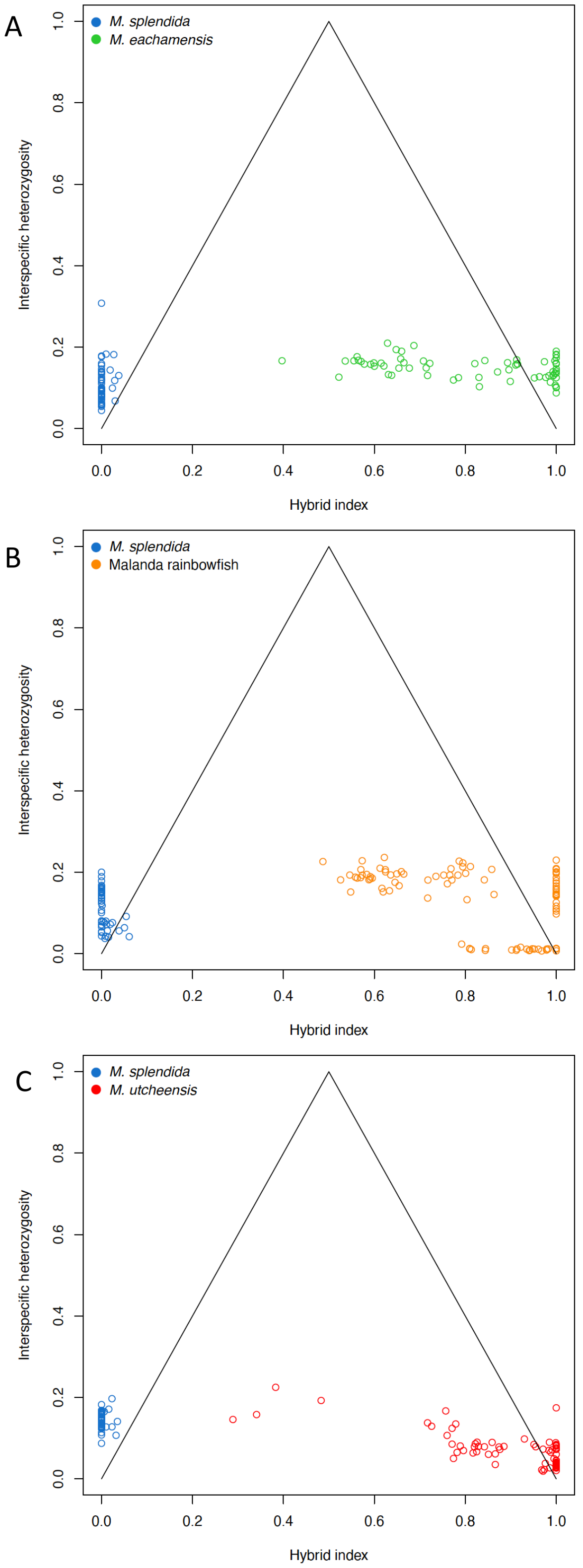


**Extended data Figure 2. Triangle plots contrasting hybrid index with interspecific heterozygosity among *Melanotaenia splendida* and A) *M.eachamensis*, B) Malanda rainbowfish and C) *M. utcheensis* populations.** Parental interspecific heterozygosity is marginally greater than expected for all species, suggesting the presence of ancestral polymorphisms in both parental species. Hybrid individuals show reduced levels of interspecific heterozygosity, providing evidence for advanced-generation hybrids.

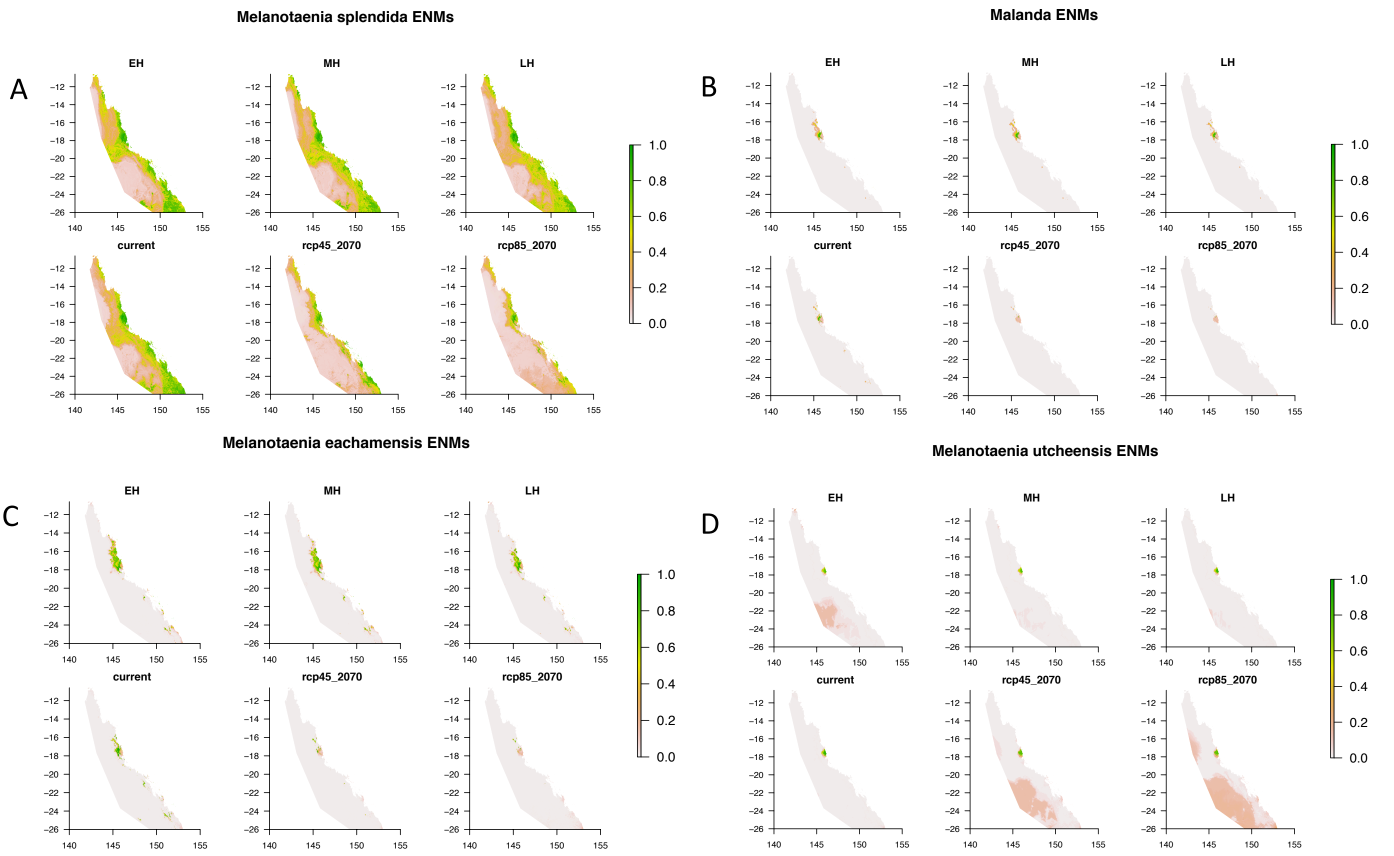


**Extended data Figure 3. Weighted mean of probabilities ensemble environmental niche models for A) *Melanotaenia splendida*, B) Malanda rainbowfish, C) *M. eachamensis*, and D) *M. utcheensis* built from individual Maximum Entropy, Generalised Linear Model, Generalised Boosting Model, and Random Forest models.** Climate models for the early-Holocene (11.7-8.326 ka), mid-Holocene (8.326-4.2 ka) and late-Holocene (4.2-0.3 ka) and projected models for 2070 under intermediate (RCP4.5) and high (RCP8.5) emissions scenarios were obtained from CHELSA v1.1 and based on the Australian Community Climate and Earth System Simulator (ACCESS1.0) global circulation model. Plot axes are (°E, °S).

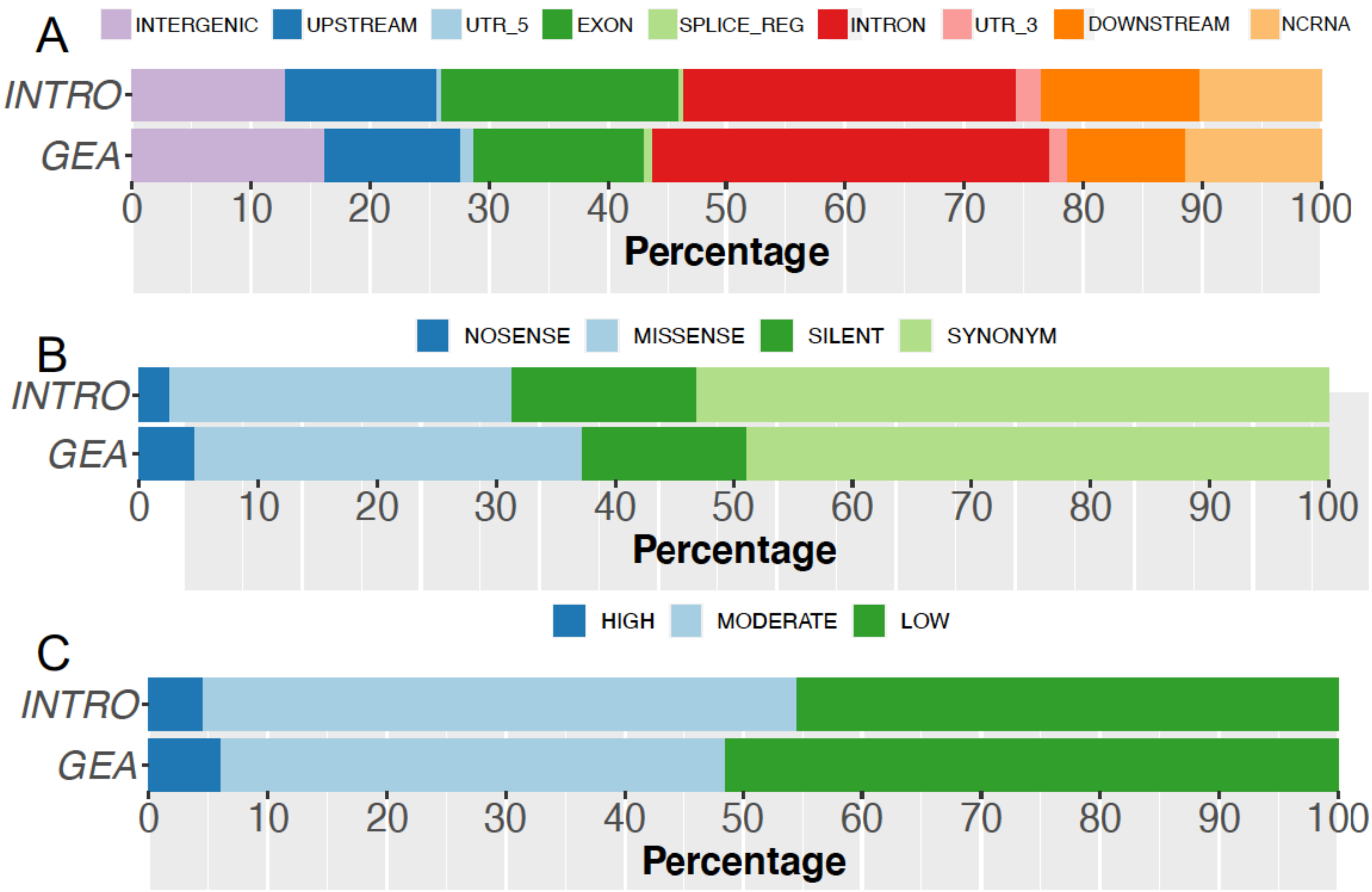


**Extended data Figure 4.** **Annotation by A) genomic region, B) functional effect, and C) impact of 211 genome environment association (GEA) candidates, and 301 introgression regions (INTRO) candidates, based on the crimson spotted rainbowfish (*Melanotaenia duboulayi*) genome annotation.**
