## Supplemental material for "Natural hybridisation reduces vulnerability to climate change"

**Supplementary Table 1. Sampling locations for rainbowfish from the Wet Tropics of Queensland, Australia.**

| Species | Site | X | Y | Museum code | Location |
|---|---|---|---|---|---|
| M. splendida | 01 | 145.643 | -17.363 | PU15-65 | North Johnstone River, Glen Allyn Road |
| M. splendida | 02 | 145.655 | -17.410 | PU16-110 | Unnamed creek on Wallace Road, below lowest falls |
| Malanda | 03 | 145.593 | -17.373 | PU16-87 | Lower Williams Creek east branch at Millaa-Millaa Road just south of Malanda |
| Malanda | 04 | 145.583 | -17.387 | PU16-89 | Tributary to Williams Creek west branch at southern end of Thiaki Creek Road |
| Malanda | 05 | 145.589 | -17.377 | PU16-90 | Tributary to Williams Creek west branch at northern end of Thiaki Creek Road |
| Malanda | 06 | 145.573 | -17.400 | PU16-131 | Mid Williams Creek west branch downstream of Timmins Road immediately below small dam |
| Malanda | 07 | 145.659 | -17.395 | PU16-100 | Unnamed creek on Wallace Road, below upper falls |
| Malanda | 08 | 145.658 | -17.396 | PU16-101 | Brodie Creek on Glen Allyn Road, just on edge of tea plantation |
| Malanda | 09 | 145.597 | -17.395 | PU16-88 | Upper Williams Creek east branch at Millaa-Millaa Road at Jaggan |
| Malanda | 10 | 145.571 | -17.400 | PU16-86 | Upper Williams Creek west branch off Thimmins Road |
| M. splendida | 11 | 145.600 | -16.807 | PU15-55 | Barron River at mouth of Owen Creek, off Oak Forest Road, west of Myola |
| M. splendida | 12 | 145.501 | -17.080 | PU16-139 | Douglas Creek on Tinaroo Creek Road |
| M. eachamensis | 13 | 145.476 | -17.262 | PU15-60 | Prior Creek at Kennedy Highway, Atherton |
| M. eachamensis | 14 | 145.491 | -17.390 | PU16-121 | Poona Creek above Falls north of Plath Road crossing, ~14 km south of Atherton |
| M. eachamensis | 15 | 145.496 | -17.384 | PU16-122 | Upper Barron River at Plath Road, ~13 km south of Atherton |
| M. eachamensis | 16 | 145.485 | -17.349 | PU16-124 | Gowrie Creek at Deep Creek Road, ~9 km south of Atherton |
| M. splendida | 17 | 145.558 | -17.336 | PU16-92 | Nicholas Creek at Andrickson Road |
| M. eachamensis | 18 | 145.600 | -17.451 | PU15-66 | Dirran Creek at Millaa Millaa-Malanda Road |
| M. eachamensis | 19 | 145.538 | -17.341 | PU16-93 | Gwynne Creek at Cook Road |
| M. eachamensis | 20 | 145.526 | -17.099 | PU17-19 | Emu Creek on Tinaroo Creek Road |
| M. eachamensis | 21 | 145.492 | -17.354 | PU17-20 | Spider Creek at Deep Creek Road, ~10 km south of Atherton |
| Malanda | 22 | 145.658 | -17.396 | PU17-21 | Unnamed creek on Wallace Road, below upper falls |
| M. utcheensis | 23 | 145.892 | -17.560 | PU15-72 | Berners Creek at Nerada Road |
| M. utcheensis | 24 | 145.912 | -17.522 | KM082 | Waraker Creek |
| M. utcheensis | 25 | 145.960 | -17.520 | KM093 | Berner Ck |
| M. utcheensis | 26 | 145.907 | -17.565 | KM086 | Fisher Ck at Fisher Ck Rd |
| M. utcheensis | 27 | 145.872 | -17.593 | KM096 | Fisher Ck above Gregory Falls |
| M. utcheensis | 28 | 145.862 | -17.550 | KM092 | Rankin Ck at Pullom Rd |
| M. utcheensis | 29 | 145.842 | -17.552 | KM091 | Bora Ck upper |
| M. utcheensis | 30 | 145.962 | -17.628 | KM083 | Utchee Ck, plunge pool below Canefield Falls |
| M. utcheensis | 31 | 145.940 | -17.639 | KM085 | Utchee Creek at Utchee Cr rd bridge |
| M. utcheensis | 32 | 145.922 | -17.642 | PU17-30 | Utchee Creek at Beahr Road |
| M. utcheensis | 33 | 145.962 | -17.666 | KM023 | Miskin Creek at Innisfail-Jappoon road crossing |
| M. utcheensis | 34 | 145.974 | -17.677 | KM084 | Meuanbah Creek |
| M. splendida | 35 | 145.926 | -18.005 | PU15-76 | Weiss Creek at Davidson Road, Euramo |
| Tully | 36 | 145.564 | -17.826 | PU15-69 | Nitchaga Creek at Tully Falls Road |
| Tully | 37 | 145.612 | -17.528 | PU17-29 | North Beatrice River at Old Palmerston Highway, south of Millaa Millaa |
| M. eachamensis | 38 | 145.615 | -17.598 | PU16-118 | Maalan River at Malaan Road |

**Supplementary Table 2. Distribution of SNPs mapping to each pseudo-chromosome.** Total number of SNPs (SNP), mean number of base pairs per SNP (SNP rate), number of genome-environment association candidate SNPs (GEA), mean number of base pairs per GEA SNP (GEA rate), number of candidate introgressed SNPs (INTRO), mean number of base pairs per INTRO SNP (INTRO rate).

| Chromosome | Length (bp) | SNP | SNP rate | GEA | GEA rate | INTRO | INTRO rate |
|---|---|---|---|---|---|---|---|
| chr_scaffold1 | 43,166,773 | 674 | 64,046 | 12 | 3,597,231 | 26 | 1,660,261 |
| chr_scaffold2 | 40,031,322 | 576 | 69,499 | 4 | 10,007,831 | 19 | 2,106,912 |
| chr_scaffold3 | 40,071,662 | 672 | 59,630 | 7 | 5,724,523 | 6 | 6,678,610 |
| chr_scaffold4 | 39,512,416 | 627 | 63,018 | 12 | 3,292,701 | 14 | 2,822,315 |
| chr_scaffold5 | 38,622,588 | 441 | 87,580 | 3 | 12,874,196 | 19 | 2,032,768 |
| chr_scaffold6 | 38,224,287 | 582 | 65,677 | 1 | 38,224,287 | 10 | 3,822,429 |
| chr_scaffold7 | 37,344,276 | 574 | 65,060 | 3 | 12,448,092 | 5 | 7,468,855 |
| chr_scaffold8 | 36,943,947 | 603 | 61,267 | 4 | 9,235,987 | 12 | 3,078,662 |
| chr_scaffold9 | 36,590,317 | 608 | 60,181 | 13 | 2,814,640 | 17 | 2,152,372 |
| chr_scaffold10 | 36,486,938 | 633 | 57,641 | 8 | 4,560,867 | 13 | 2,806,688 |
| chr_scaffold11 | 36,725,740 | 545 | 67,387 | 13 | 2,825,057 | 24 | 1,530,239 |
| chr_scaffold12 | 36,196,323 | 588 | 61,558 | 6 | 6,032,721 | 9 | 4,021,814 |
| chr_scaffold13 | 35,464,471 | 598 | 59,305 | 6 | 5,910,745 | 4 | 8,866,118 |
| chr_scaffold14 | 34,396,068 | 645 | 53,327 | 2 | 17,198,034 | 0 | NA |
| chr_scaffold15 | 34,170,226 | 535 | 63,870 | 8 | 4,271,278 | 2 | 17,085,113 |
| chr_scaffold16 | 33,535,967 | 566 | 59,251 | 8 | 4,191,996 | 41 | 817,950 |
| chr_scaffold17 | 33,778,492 | 566 | 59,679 | 7 | 4,825,499 | 8 | 4,222,312 |
| chr_scaffold18 | 32,535,952 | 523 | 62,210 | 14 | 2,323,997 | 1 | 32,535,952 |
| chr_scaffold19 | 30,432,280 | 537 | 56,671 | 6 | 5,072,047 | 2 | 15,216,140 |
| chr_scaffold20 | 29,464,887 | 574 | 51,333 | 6 | 4,910,815 | 22 | 1,339,313 |
| chr_scaffold21 | 28,992,765 | 478 | 60,654 | 6 | 4,832,128 | 17 | 1,705,457 |
| chr_scaffold22 | 28,223,336 | 539 | 52,362 | 8 | 3,527,917 | 4 | 7,055,834 |
| chr_scaffold23 | 25,807,574 | 465 | 55,500 | 12 | 2,150,631 | 9 | 2,867,508 |
| chr_scaffold24 | 23,383,415 | 399 | 58,605 | 2 | 11,691,708 | 15 | 1,558,894 |

**Supplementary Table 3. Genetic diversity summary statistics for rainbowfish from the Wet Tropics of Queensland, Australia.** Species, hybrid status, site number, sample size (N), expected heterozygosity (He), observed heterozygosity (Ho) and percentage of polymorphic loci (%poly).

| Species | Status | Site | N | He | Ho | %poly |
|---|---|---|---|---|---|---|
| *M. splendida* | Pure | 01 | 12 | 0.157 | 0.144 | 57.7 |
| | Pure | 02 | 12 | 0.129 | 0.129 | 42.2 |
| | Pure | 11 | 12 | 0.135 | 0.120 | 47.0 |
| | Pure | 12 | 12 | 0.126 | 0.113 | 42.3 |
| | Pure | 17 | 10 | 0.144 | 0.131 | 43.3 |
| | Pure | 35 | 12 | 0.122 | 0.112 | 42.8 |
| Malanda | Pure | 07 | 12 | 0.008 | 0.008 | 2.5 |
| | Pure | 08 | 12 | 0.011 | 0.010 | 4.8 |
| | Pure | 09 | 12 | 0.046 | 0.045 | 21.6 |
| | Pure | 10 | 12 | 0.071 | 0.062 | 35.0 |
| | Pure | 22 | 4 | 0.091 | 0.096 | 26.7 |
| | Hybrid | 03 | 12 | 0.164 | 0.155 | 51.9 |
| | Hybrid | 04 | 12 | 0.141 | 0.140 | 46.4 |
| | Hybrid | 05 | 10 | 0.136 | 0.130 | 46.3 |
| | Hybrid | 06 | 12 | 0.111 | 0.106 | 43.0 |
| *M. eachamensis* | Pure | 16 | 12 | 0.112 | 0.107 | 41.1 |
| | Pure | 18 | 12 | 0.071 | 0.064 | 27.5 |
| | Pure | 19 | 7 | 0.080 | 0.072 | 26.0 |
| | Pure | 20 | 11 | 0.085 | 0.078 | 35.0 |
| | Pure | 38 | 10 | 0.086 | 0.081 | 33.0 |
| | Hybrid | 13 | 9 | 0.156 | 0.149 | 46.5 |
| | Hybrid | 14 | 2 | 0.124 | 0.107 | 21.7 |
| | Hybrid | 15 | 8 | 0.157 | 0.143 | 46.3 |
| | Hybrid | 21 | 8 | 0.149 | 0.131 | 44.5 |
| *M. utcheensis* | Pure | 23 | 12 | 0.049 | 0.047 | 21.9 |
| | Pure | 24 | 8 | 0.083 | 0.069 | 25.2 |
| | Pure | 25 | 2 | 0.067 | 0.080 | 13.0 |
| | Pure | 26 | 8 | 0.033 | 0.032 | 10.3 |
| | Pure | 27 | 7 | 0.039 | 0.041 | 14.5 |
| | Pure | 28 | 7 | 0.070 | 0.073 | 20.4 |
| | Pure | 29 | 8 | 0.047 | 0.046 | 19.9 |
| | Pure | 31 | 4 | 0.089 | 0.088 | 23.0 |
| | Pure | 32 | 4 | 0.075 | 0.074 | 18.7 |
| | Pure | 33 | 7 | 0.083 | 0.084 | 27.6 |
| | Pure | 34 | 5 | 0.078 | 0.082 | 21.6 |
| | Hybrid | 30 | 5 | 0.164 | 0.143 | 40.7 |
| Tully | Pure | 36 | 12 | 0.103 | 0.097 | 31.8 |
| | Pure | 37 | 6 | 0.141 | 0.117 | 39.6 |

**Supplementary Table 4. Genetic diversity summary statistics for rainbowfish from the Wet Tropics of Queensland, Australia aggregated for pure and hybrid populations of each species.** Species, hybrid status, sample size (N), expected heterozygosity (He), observed heterozygosity (Ho) and percentage of polymorphic loci (%poly).

| Species | Status | N | He | Ho | %poly |
|---|---|---|---|---|---|
| *M. splendida* | Pure | 70 | 0.136 | 0.125 | 45.9 |
| Malanda | Pure | 52 | 0.045 | 0.044 | 18.1 |
| | Hybrid | 46 | 0.138 | 0.133 | 46.9 |
| *M. eachamensis* | Pure | 52 | 0.087 | 0.080 | 32.5 |
| | Hybrid | 27 | 0.147 | 0.133 | 39.7 |
| *M. utcheensis* | Pure | 72 | 0.065 | 0.065 | 19.6 |
| | Hybrid | 5 | 0.164 | 0.143 | 40.7 |
| Tully | Pure | 18 | 0.122 | 0.107 | 35.7 |

**Supplementary Table 5. NewHybrids classifications for *Melanotaenia splendida* x narrow endemic species (NER).** Possible hybrid classes are first generation crosses NER x *M. splendida* (F1), F1 x F1 crosses (F2), NER x F1 back-crosses (BC1), and *M. splendida* x F1 back-crosses (BC2).

| NER species | F1 | F2 | BC1 | BC2 |
|---|---|---|---|---|
| *M. eachamensis* | 0 | 14 | 13 | 0 |
| Malanda | 0 | 25 | 22 | 0 |
| *M.utcheensis* | 0 | 2 | 1 | 2 |

**Supplementary Table 6. Model evaluation for the weighted mean of probabilities ensemble environmental niche models for each species.**

| Species | Metric | Eval | Cutoff | Sensitivity | Specificity |
|---|---|---|---|---|---|
| *M. splendida* | TSS | 0.622 | 443.5 | 89.55 | 72.06 |
| | ROC | 0.898 | 460.5 | 88.25 | 74.15 |
| *M. eachamensis* | TSS | 0.987 | 349.0 | 100.00 | 98.73 |
| | ROC | 0.999 | 354.0 | 100.00 | 98.80 |
| Malanda | TSS | 1.000 | 743.0 | 100.00 | 100.00 |
| | ROC | 1.000 | 747.0 | 100.00 | 100.00 |
| *M. utcheensis* | TSS | 1.000 | 858.0 | 100.00 | 100.00 |
| | ROC | 1.000 | 859.5 | 100.00 | 100.00 |

**Supplementary Table 7. Results for the partial redundancy analysis.**

| | Inertia | Proportion | Rank |
|---|---|---|---|
| Total | 744.927 | 1.0000 | |
| Conditional | 219.949 | 0.2953 | 3 |
| Constrained | 110.5069 | 0.1483 | 2 |
| Unconstrained | 414.4711 | 0.5564 | 32 |

**Supplementary Table 8. Results for the permutation test on the redundancy analysis.**

| | Df | Variance | F | Pr(>F) |
|---|---|---|---|---|
| Model | 2 | 110.51 | 4.2659 | 0.001 |
| Residual | 32 | 414.47 | | |
